# Maximizing cooperation in the prisoner’s dilemma evolutionary game via optimal control

**DOI:** 10.1101/2020.07.13.201400

**Authors:** P.K. Newton, Y. Ma

## Abstract

The prisoner’s dilemma (PD) game offers a simple paradigm of competition between two players who can either cooperate or defect. Since defection is a strict Nash equilibrium, it is an asymptotically stable state of the replicator dynamical system that uses the PD payoff matrix to define the fitness landscape of two interacting evolving populations. The dilemma arises from the fact that the average payoff of this asymptotically stable state is sub-optimal. Coaxing the players to cooperate would result in a higher payoff for both. Here we develop an optimal control theory for the prisoner’s dilemma evolutionary game in order to maximize cooperation (minimize the defector population) over a given cycle-time *T*, subject to constraints. Our two time-dependent controllers are applied to the off-diagonal elements of the payoff matrix in a bang-bang sequence that dynamically changes the game being played by dynamically adjusting the payoffs, with optimal timing that depends on the initial population distributions. Over multiple cycles *nT* (*n* > 1), the method is adaptive as it uses the defector population at the end of the *n^th^* cycle to calculate the optimal schedule over the *n* + 1^*st*^ cycle. The control method, based on Pontryagin’s maximum principle, can be viewed as determining the optimal way to dynamically alter incentives and penalties in order to maximize the probability of cooperation in settings that track dynamic changes in the frequency of strategists, with potential applications in evolutionary biology, economics, theoretical ecology, and other fields where the replicator system is used.

**PACS numbers:** 02.50.Le; 02.30.Yy; 05.45.-a; 87.23.Kg; 87.23.Cc

## I. INTRODUCTION

Game theory models, originally developed by von Neumann and Morgenstern in 1944 [1], are widely used as paradigms to study cooperation and conflict in fields ranging from military strategy [2], social interactions [3], economics [1], computer science [4], the physics of complex systems [5], evolutionary psychology [6], evolutionary biology [7], and more recently, cancer [8–12]. To characterize the game, a payoff matrix, *A*, is introduced which for a two player game is of the generic form:

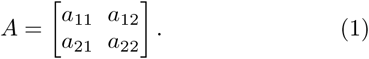

The entries of the payoff matrix determine the costbenefit trade-off associated with each game between the two competitors who rationally compete with the goal of maximizing their own payoff [13]. We show in figure 1 12 possible games that can be played (out of a total of 78 [14]) using (1) as the payoff matrix. Notice for a given value of *a*_22_ which we place at the origin without loss of generality, and with an *a*_11_ > *a*_22_ (hence the reduction in the total number of games), we can play any game if we choose the off-diagonal elements of A appropriately in the (*a*_12_, *a*_21_) plane. The prisoner’s dilemma (PD) region of the plane occupies special territory among all other regions as it is by far the most studied and used in models that focus on the evolution of cooperation [15–17]. A well studied framework is the iterated PD game between two players that are allowed to both view their opponent’s past strategies, and adjust their own strategy for each new game based on past information. Predicting which strategy will work best is difficult, and the famous Axelrod tournaments in which competitors submitted their strategy, and the pool of strategies competed through computer simulations to see which ones worked best was the beginning of the study of such systems [3]. Recent contributions to this literature introduced a new, previously undiscovered successful strategy [18].

**FIG. 1.**
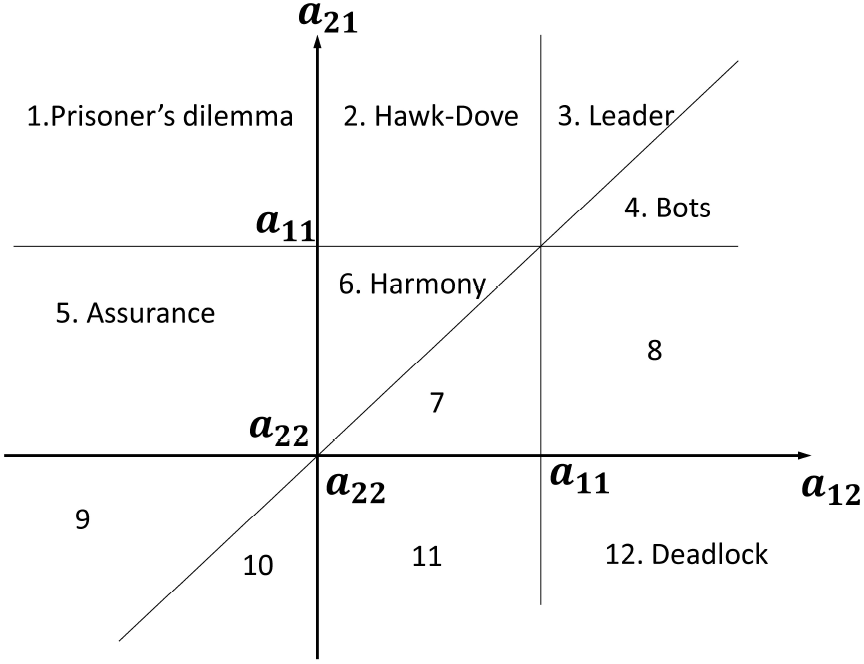
Twelve regions in the (*a*_12_, *a*_21_) plane define which of the games is being played. There is no loss of generality in choosing *a*_22_ at the origin.

Evolutionary game theory, used in population dynamics models involving evolution by natural selection [7] similarly, makes use of the payoff matrix (1), but embeds it directly into a dynamical setting by assigning the payoff matrix to a dynamical system and associates payoff with reproductive prowess. A common evolutionary game theory model is the replicator dynamical system [19], in the context of two population frequencies 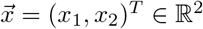:

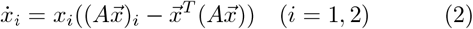

with *x*_1_ + *x*_2_ = 1, 0 ≤ *x*_1_ ≤ 1,0 ≤ *x*_2_ ≤ 1, where each variable has the alternative interpretation as a probability. One can also think of the variables 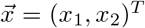 as strategies that evolve, with the most successful strategy, say *x*_2_(*t*) → 1, while the other *x*_1_(*t*) → 0 (as in the PD game). 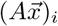 is the fitness of population *i*, and 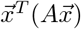 is the average fitness of both populations, so the term multiplying *x_i_* in (2) drives growth if the population *i* is above the average and decay if it is below the average. The fitness functions in (2) are said to be population dependent (selection pressure is imposed by the mix of population frequencies) and determine growth or decay of each subpopulation.

The PD game captures, in a simple framework, the competition between two players, one of whom is labeled a cooperator (say *x*_1_), while the other is labeled a defector (say *x*_2_). As such, the prisoner’s dilemma game presents a situation where if each player cooperated, they would receive a better payoff than if they both defected. The dilemma is, if they are rational players, they will both choose to defect, as this state is a Nash equilibrium of the system [20], defined as a strategy 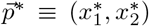 where 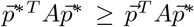, for all 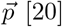 [20]. This paradigm is well understood and has been discussed extensively in the literature [15, 16].

Motivated by the use of the replicator equations and PD payoff matrix in cancer models [21, 22], we develop an optimal control framework for the replicator equations (2) by allowing the off-diagonal elements of the payoff matrix, *a*_12_(*t*), *a*_21_(*t*), to be time-dependent functions that allow us to exogenously shape the fitness landscape of two co-evolving populations. As these fitness values change in time, we move to different regions of the plane shown in figure 1 which dynamically changes the *instantaneous* game being played during the course of the population evolution. This interpretation has been suggested recently in the context of developing adaptive therapy schedules for cancer [11, 12]. The interplay between the timescales associated with the control functions *a*_12_(*t*) and *a*_21_(*t*) and the underlying dynamical timescales associated with the replicator system (2) makes the optimal control problem interesting. As far as we are aware, the only works we know of in which the payoff matrix is altered during the course of evolution in the context of the replicator equations is the interesting paper by Weitz et al [23] who allow the payoff entries to co-evolve, using feedback, along with the populations. Our goal in this paper is to show how to optimally reduce the defector population *x*_2_ (i.e. maximize cooperation) after a fixed cycle-time *T* in which we apply our time-dependent controller schedule. Then we extend the method to *n* (up to *n* = 5) cycle times *nT* with an adaptive method that uses the defector population at the end of the *n^th^* cycle, *x*_2_(*nT*), to compute the optimal control schedule for the *n*+1^*st*^ cycle and *x*_2_((*n* + 1)*T*). A nice recent review paper on the combined use of game theory and control is [24].

## II. REPLICATOR DYNAMICS AND OPTIMAL CONTROL THEORY

### A. Replicator dynamical system

We start with the uncontrolled payoff matrix of prisoner’s dilemma type:

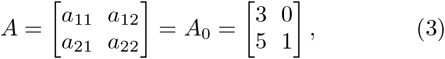

where the population *x*_1_ are the cooperators, and *x*_2_ are the defectors. The Nash equilibrium, 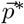, is the defector state *x*_1_ = 0, *x*_2_ = 1, which is easily shown to be a strict Nash equilibrium since 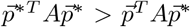 for all 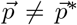. This implies that the defector state is also an evolutionary stable state (ESS) of the replicator system (2) as discussed in [20].

We then introduce an augmented payoff matrix *A* of the form:

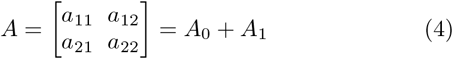

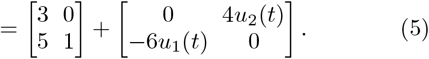

where *A*_1_ represents our control. The time-dependent functions 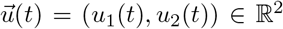 are bounded above and below, 0 ≤ *u*_1_(*t*) ≤ 1, 0 ≤ *u*_2_(*t*) ≤ 1 and have a range that allows use to traverse the plane depicted in figure 1 to play each of the 12 possible games at any given snapshot in time. We will use the two control functions to *shape the fitness landscape* of the system appropriately by dynamically changing the game being played between the two populations as the system evolves (only the difference in the row values of *A*_0_ determine the dynamics which makes two controllers sufficient).

To motivate our problem further and understand why the prisoner’s dilemma game has been used as a paradigm for tumor growth [9, 21, 22, 25], think of a competing population of healthy cells, *x*_1_, and cancer cells, *x*_2_, where the healthy cells play the role of cooperators and the cancer cells play the role of the defectors in a PD evolutionary game [8]. Using (2), starting with any tumor cell population 0 < *x*_2_ ≤ 1, it is straightforward to show: (i) *x*_2_ → 1, as *t* → ∞, (ii) the average fitness of the healthy cell state *x*_1_ = 1, *x*_2_ = 0 is greater than the average fitness of the cancer cell state *x*_1_ = 0, *x*_2_ = 1 since 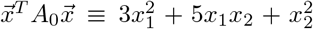 is the average fitness and 3 > 1, (iii) the average fitness of the total population decreases as the cancer cell population saturates, and (iv) the tumor growth curve *x*_2_(*t*) vs. *t* yields an S-shaped curve very typical of tumor growth curves [26].

In this context, the controller 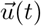 can be thought of as a chemotherapy schedule which will alter the fitness balance of the uncontrolled healthy cell/tumor cell populations. The goal of a simple chemotherapy schedule might be to shape the fitness landscape so that the defector (tumor) cells *x*_2_(*t*) do not reach the saturated state *x*_2_(*t*) = 1. If we denote the dose 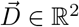:

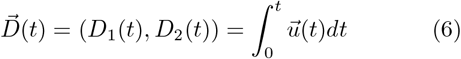

as the total dose delivered in time *t*, then:

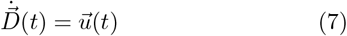

and:

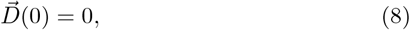

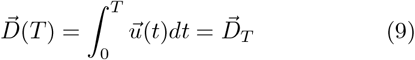

where *T* denotes a final time in which we implement the control. Thus, 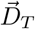 denotes the total dose delivered in time *T* which we will use that as a constraint on the optimization problem. Then, our control goal is to maximally reduce the tumor cell (defector) population *x*_2_(*t*) at the end of one chemotherapy cycle 0 ≤ *t* ≤ *T*, given constraints on the total dose delivered. We are particularly interested in the optimal schedule 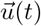 that produces the optimal response and we will compare it with the responses produced by other schedules (that have chemotherapeutic interpretations). Stated simply, we are interested in obtaining the time-dependent schedule (*u*_1_(*t*), *u*_2_(*t*)) in (5) that maximizes cooperation *x*_1_(*t*) for the system (2) at the end of time *t* = *T*. This general framework extends to most other applications in which evolutionary game theory is used.

### B. Pontryagin maximum principle

We utilize the standard form for implementing the maximum (minimum) principle with boundary value con-straints:

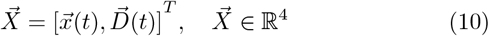

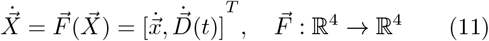

with the goal of minimizing a general cost function [27–30]:

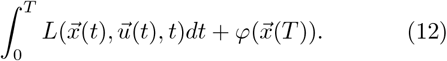

Following [30], we construct the control theory Hamiltonian:

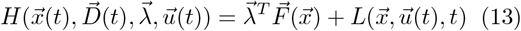

where 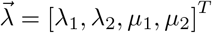 are the co-state functions (i.e. momenta) associated with 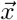 and 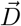 respectively. Assum-ing that 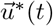 is the optimal control for this problem, with corresponding trajectory 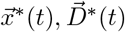, the canonical equations satisfy:

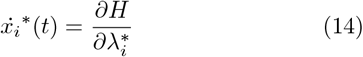

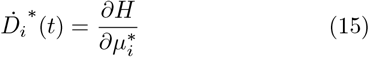

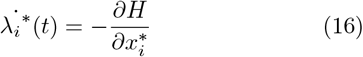

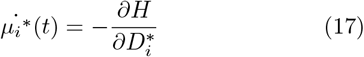

where *i* = (1, 2). The corresponding boundary conditions are:

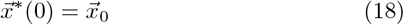

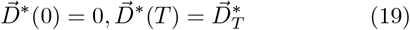

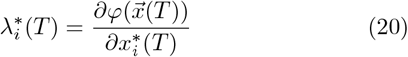

Then, at any point in time, the optimal control 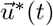 will minimize the control theory Hamiltonian:

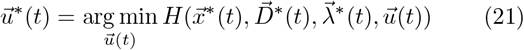

The optimization problem becomes a two-point boundary value problem (using (18)-(20)) with unknowns 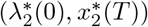 whose solution gives rise to the optimal trajectory 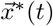 (from (14)) and the corresponding control 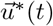 that produces it [27–30]. For the optimization, we take 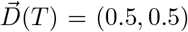 and to minimize the defector frequency at the end of one cycle *t* = *T*, we choose our cost function (12):

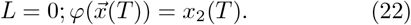

We solve this problem by standard shooting type methods [31].

## III. RESULTS

Because the Hamiltonian (13) is linear in the controllers (with *L* = 0 in (21)), it is straightforward to prove that the optimal control 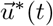 is bang-bang. We compare the optimal schedule and trajectories with two other (non-optimal) ones in figure 2, for a small value of *x*_2_(0) =0.1 (figure 2(a),(b)), intermediate value of *x*_2_(0) = 0.5 (figure 2(c),(d)), large value of *x*_2_(0) = 0.9 (figure 2(e),(f)), and a value near saturation *x*_2_(0) = 0.95 (figure 2(g),(h)). There are several interesting points to make. First, for initial values *x*_2_(0) < 0.95, the optimal control schedule allows the defector proportion to increase for a short time before *u*_2_ turns on, which is perhaps counter intuitive. For initial values sufficiently small, this initial growth phase is compensated by keeping *u*_1_ turned on until the end of the cycle time *T* (figures 2(b),(d)). The larger the initial defector proportion, the earlier the control *u*_2_ turns on (figures 2(f),(h)), until for values above a threshold of *x*_2_(0) ~ 0.91, the initial time abruptly goes to zero (figure 3(b)). It is more beneficial to control the growth of *x*_2_(*t*) at the beginning of the cycle than at the end (figure 2(g),(h)). Also, notice that for the optimal control sequences shown in figures 2(b),(d),(f),(h), the controllers *u*_1_ and *u*_2_ partially overlap in the time they are turned on in the middle of the cycle at the expense of leaving both off either at the beginning or end. We compare the optimal trajectories (figures 2(a),(c),(e)) with the case where there is no time-overlap (i.e. sequential: first *u*_2_ followed by *u*_1_), and the case where *u*_1_ and *u*_2_ are held constant throughout the cycle time *T*, all with the same value of 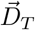. For small initial values (figure 2(a),(b)) there is very little difference between the optimal trajectory and the one produced by a sequential controller, particularly for small initial con-ditions.

**FIG. 2.**
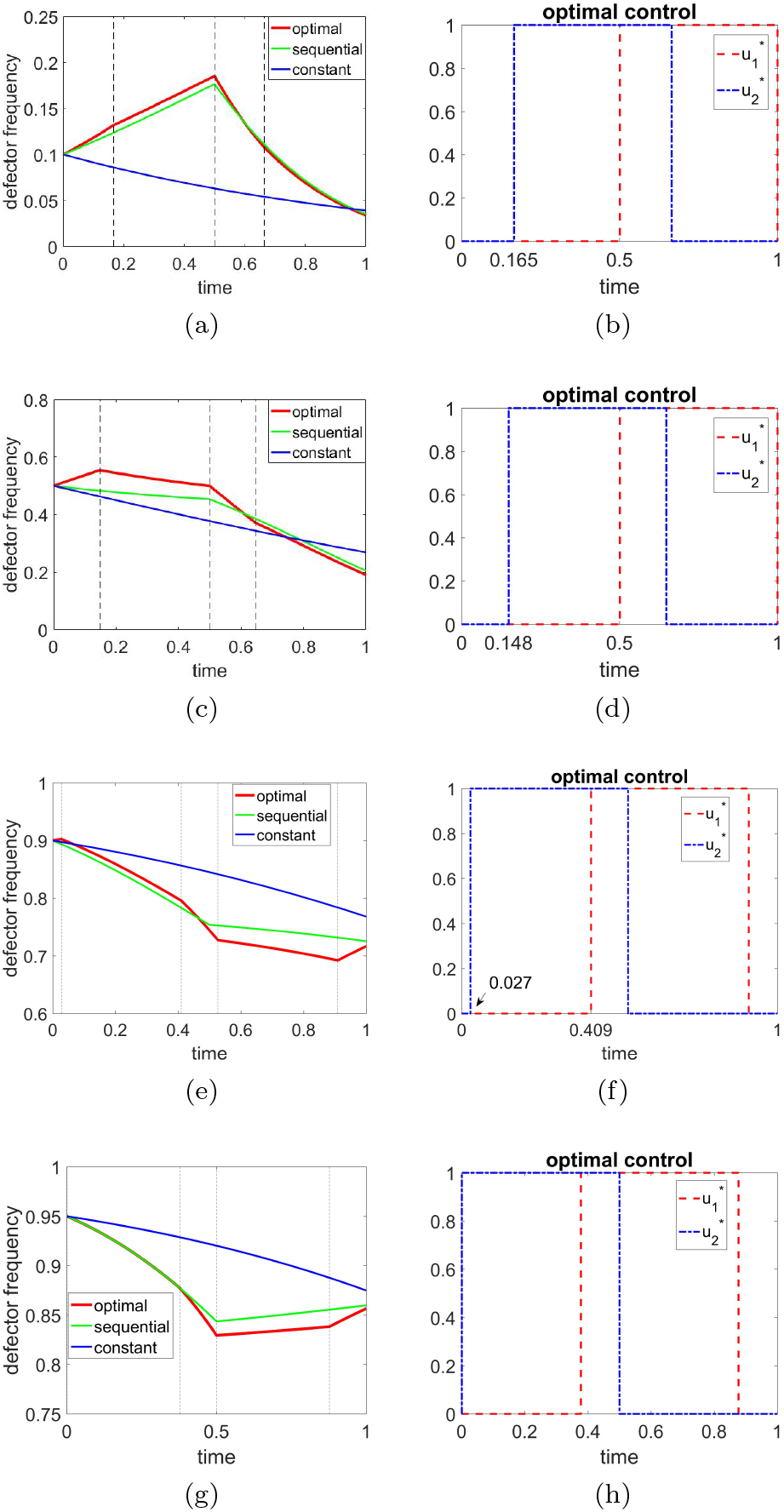
Optimal trajectories and optimal control sequences for small, medium, and large initial defector populations, with 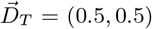. (a) Defector frequency for initial state *x*_2_(0) = 0.1; (b) Optimal control sequence for initial state *x*_2_(0) = 0.1; (c) Defector frequency for initial state *x*_2_(0) = 0.5; (d) Optimal control sequence for initial state *x*_2_(0) = 0.5; (e) Defector frequency for initial state *x*_2_(0) = 0.9; (f) Optimal control sequence for initial state *x*_2_(0) = 0.9;(g) Defector frequency for initial state *x*_2_(0) = 0.95; (h) Optimal control sequence for initial state *x*_2_(0) = 0.95.

In figure 3(a) we show the percentage reduction of the defector frequency (1 – *x*_2_(T)/*x*_2_(0)) as a function of the initial frequency *x*_2_(0). The maximum reduction (dashed vertical line) occurs for an initial defector proportion of around 22%. The curve also shows that the optimal schedule is much more efficient at reducing the proportion of defectors for small populations than for large ones, with reduction approaching zero as the initial defector population approaches one. Also, in the limits *x*_2_(0) → 0,1, all three schedules converge to the same value, showing that the exact schedule 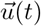 matters less than the total dose 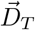 which is the same for each. This is because in those regimes the system is essentially linear, and the exact solution depends only on 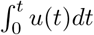. Figure 3(b) shows the time at which each of the controllers is turned on, with an abrupt change for large enough initial defector populations *x*_2_(0) ~ 0.91 as mentioned.

**FIG. 3.**
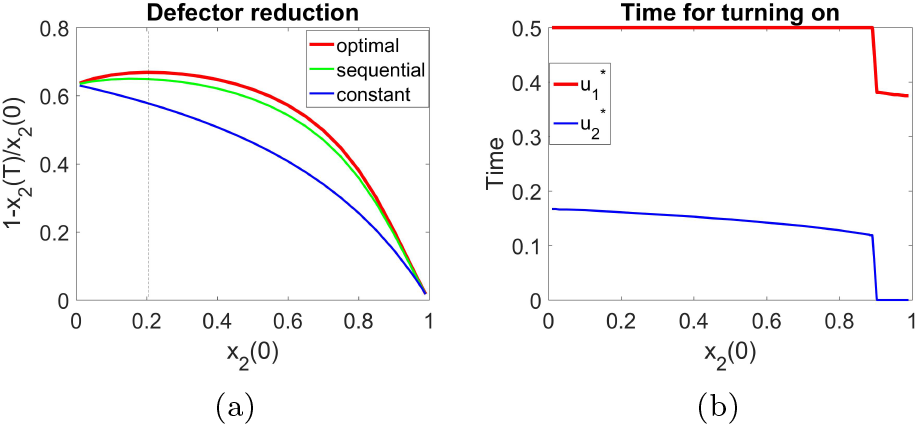
(a) Percentage reduction as a function of initial defector proportion. Maximum reduction (shown as dashed vertical line) occurs for initial defector proportion of around 22%. In the limits *x*_2_(0) → 0,1, the exact schedule 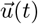 does not matter, only the total dose 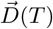. The controls schedule is much more efficient at reducing the proportion of defectors for small populations than for large; (b) Turn-on time for *u*_1_(*t*) and *u*_2_(*t*) as a function of initial defector population *x*_2_(0). In the narrow (approximate) range 0.89 ≤ *x*_2_(0) ≤ 0.91, the times change abruptly and the control sequence has 5 segments as shown in figure 2(e),(f).

Figure 4 shows the sequence of distinct games that we cycle through to produce the optimal trajectory:

1. **Prisoner’s dilemma** (asymptotically stable state *x*_2_ = 1);
2. **Leader** (asymptotically stable state *x*_2_ = 2/5);
3. **Deadlock** (asymptotically stable state *x*_2_ = 0);
4. **Game** #10 in figure 1 (asymptotically stable states *x*_2_ = 0, 1).

**FIG. 4.**
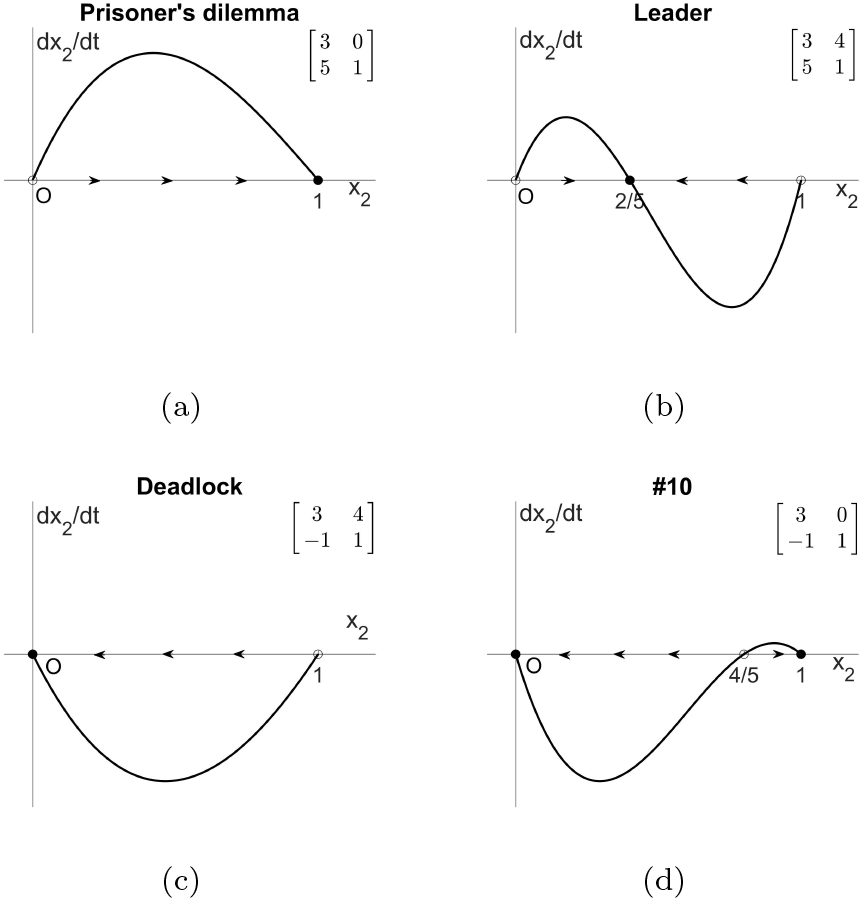
The sequence of four games that produce the optimal trajectory if the switching times are chosen as shown in figure 2. (a) The first game is a prisoner’s dilemma game, where *x*_2_ = 1 is an asymptotically stable fixed point; (b) The second game is a Leader game with interior fixed point *x*_2_ = 2/5 being an asymptotically stable fixed point; (c) The third game is a Deadlock game with *x*_2_ = 0 being the asymptotically stable fixed point; (d) The fourth game is game #10 in figure 1 with *x*_2_ = 0, 1 being asymptotically stable fixed points.

This is easiest to understand by decoupling the replicator system (2) and writing the cubic nonlinear ordinary differential equation for *x*_2_:

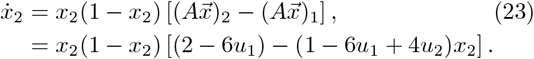

The sequence of games is obtained using:

i. *u*_1_ = 0, *u*_2_ = 0
ii. *u*_1_ = 0, *u*_2_ = 1
iii. _u1_ = 1, *u*_2_ = 1
iv. *u*_1_ = 1, *u*_2_ = 0

To produce the optimal trajectory, the switching times must be chosen as shown in figures 2(b),(d),(f), (h) (note in (f) and (h) the game switches back to PD just before *T*), and these times depend on the initial state *x*_2_(0). The flow associated with the sequence of four games depicted in figure 4 makes it clear why the switching times are tied to the initial condition.

Figure 5 shows the defector frequencies 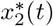 over a sequence of five cycles 0 ≤ *t* ≤ 5*T* for the optimal control sequence, as compared with the sequential and constant controllers. In the case of optimization over multiple cycles, the method is adaptive as it must use the frequency obtained at the end of the *n^th^* cycle, 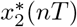, to calculate the optimal schedule associated with the *n*+1^*st*^ cycle. For large initial values *x*_2_(0) = 0.9 (shown in figures 5(a),(b)), the frequency reduction curves start decreasing (nearly) linearly, but deviates from linear over time. For small initial values (shown in figures 5(e),(f) for *x*_2_(0) = 0.1), the reduction is very close to linear.

**FIG. 5.**
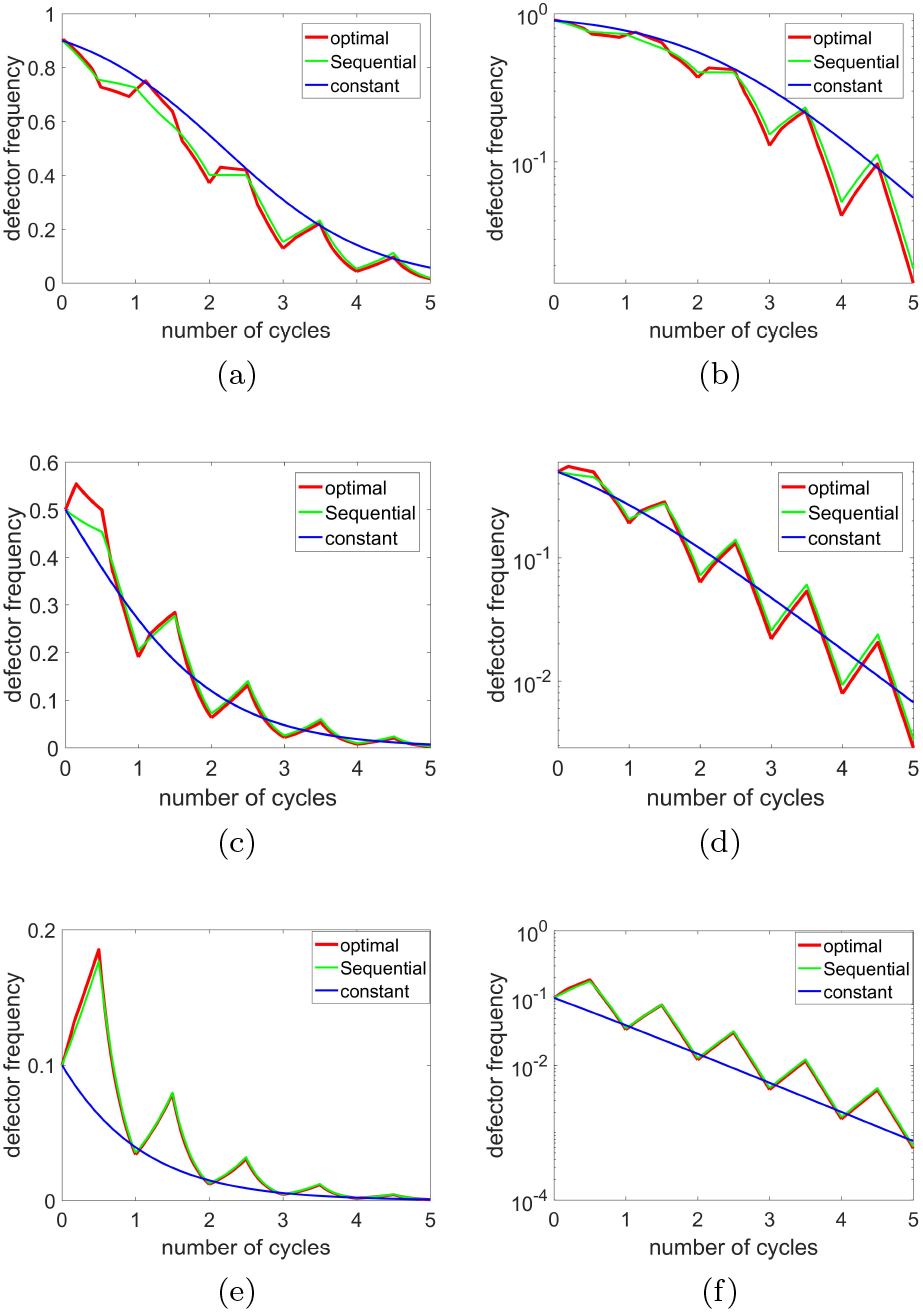
Defector frequency reduction using optimal control over five consecutive cycles.(a) Defector optimal frequency with *x*_2_(0) = 0.9 over five cycles as compared with frequency associated with sequential control and constant control; (b) Same as (a) plotted on log-linear scale; (c) Defector optimal frequency with *x*_2_(0) = 0.5 over five cycles as compared with frequency associated with sequential control and constant control; (d) Same as (c) plotted on log-linear scale; (e) Defector optimal frequency with *x*_2_(0) = 0.1 over five cycles as compared with frequency associated with sequential control and constant control; (f) Same as (e) plotted on log-linear scale.

## IV. DISCUSSION

We develop an optimal control theory for maximizing cooperation in the prisoner’s dilemma evolutionary game by dynamically altering the payoffs during the course of evolution, subject to certain constraints. The optimal control schedule is bang-bang for each of the two controllers, with switching times that depend on the initial proportion of cooperators to defectors. One interpretation of the method is that we cycle through a sequence of four distinct payoff matrix types (PD → Leader → Deadlock → #10), switching the game at precisely the right times to maximize group cooperation. Interestingly, a recent paper interprets a strategy used by non-small cell lung cancer co-cultures as one of switching from playing a Leader game to playing a Deadlock game in order to develop resistance [12].

The interpretation of the control functions depends on the specific application with many potential interpretations in the context of controlling the balance of evolving populations. Aside from optimizing chemotherapy schedules to control tumors [21, 22, 32–34], one might think of the application of antibiotic schedules to control microbial populations [35], or the strategic application of toxins to control infestations using integrated pest management protocols [36]. In economics, one might think of time-dependent payoffs as dynamic policy actions that seek to optimize certain societal economic goals [37], or in traffic flow applications, one might seek dynamic incentives for minimizing transit times by dynamic payoff schedules [38]. We see no particular technical roadblock to generalizing the optimal control method to *N* × *N* evolutionary games, although the numerical challenges of solving the necessary two-point boundary value problems becomes more severe, particularly searching for appropriate initial guesses to ensure convergence.

As a final note, we mention that a system related to (2) is the adjusted replicator equation [25] with growth/decay terms normalized by the average fitness. While the normalization term has no effect on ratios 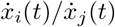, it does effect the time-scaling of each variable, hence changes the optimal control problem. Most importantly, the control Hamiltonian is no longer linear in 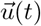, so the optimal control may not necessarily be bang-bang, and we have preliminary results that show this. Since the adjusted replicator system is known to be the deterministic limit of the finite-population stochastic Moran process in the limit of infinite populations [39, 40], this deterministic optimal control problem should be related to the stochastic optimal control problem associated with the Moran process via a Markov chain approach.

## ACKNOWLEDGMENTS

We gratefully acknowledge support from the Army Research Office MURI Award #W911NF1910269 (20192024).

